# IGF2BP2 Promotes Cancer Progression by Degrading the RNA Transcript Encoding a v-ATPase Subunit

**DOI:** 10.1101/2021.04.01.438101

**Authors:** A. Latifkar, F. Wang, J.J. Mullmann, I.R. Fernandez, L. Ling, C. Fischbach, R.S. Weiss, H. Lin, R.A. Cerione, M.A. Antonyak

## Abstract

Insulin-like growth factor 2 mRNA binding protein 2 (IGF2BP2) binds to various RNA transcripts and promotes cancer progression, although little is known regarding its regulation. Here we show IGF2BP2 is a substrate of the deacetylase and tumor suppressor sirtuin 1 (SIRT1) and regulates the expression of the vacuolar ATPase subunit ATP6V1A. SIRT1 down-regulation in aggressive cancers leads to increased acetylation of IGF2BP2 which recruits the XRN2 nuclease to degrade the ATP6V1A transcript, decreasing its expression. This impairs lysosomal function and results in the production of a secretome that enhances cancer cell proliferation and metastasis. These findings describe a previously unrecognized role for IGF2BP2 in the degradation of an mRNA transcript essential for lysosomal function and highlight how its sirtuin-regulated acetylation state can have significant biological and disease consequences.

**One Sentence Summary:** Acetylation of the RNA binding protein IGF2BP2, upon down-regulation of SIRT1, leads to degradation of the transcript encoding ATP6V1A and impaired lysosomal function in aggressive cancer cells.

## Main Text

IGF2BP2, also known as IMP2, is a member of the family of IGFBPs that are highly conserved in vertebrates (1). It was first identified as an RNA-binding protein that associates with *IGF2* mRNA and subsequently shown to bind to other transcripts including those encoding various mitochondrial proteins (1-3). The regulation and biological roles of IGF2BP2 are still being determined, although it has been suggested to be a tumor promoter based on its ability to stabilize mRNA transcripts as well as N6-methyladenosine-modified long non-coding RNAs (1-8). Building on our efforts to understand the underlying mechanism by which SIRT1 functions as a tumor suppressor (9-11), we have now uncovered a regulatory mechanism by which IGF2BP2 contributes to transcript degradation with important consequences for cancer progression.

Recently, we showed that knocking-down SIRT1 in breast cancer cells results in the reduced expression of ATP6V1A, the catalytic subunit of the v-ATPase which is essential for maintaining the acidic environment of lysosomes (12). These changes led to the increased shedding of exosomes enriched in ubiquitinated protein cargo and to the secretion of soluble hydrolases (i.e. cathepsins), producing a secretome that promoted breast cancer cell survival, migration, and invasion. To establish that these effects were not unique to SIRT1 knock-down cells, we examined the MCF10A breast cancer progression series which consists of isogenic cell lines that represent different stages of breast cancer, ranging from non-tumorigenic MCF10A epithelial cells to the highly malignant and aggressive MCF10CA1A cell line (13). The lowest levels of SIRT1 and ATP6V1A expression were detected in the malignant MCF10CA1A cells (**Fig. S1A**), accompanied by an increase in the number of shed exosomes (**Fig. S1B**). We further probed the consequences of disabling SIRT1 for breast cancer progression by treating tumor-bearing MMTV-PYMT mice (14) with EX-527, a small molecule inhibitor of SIRT1 deacetylase activity (15). Treatment with the SIRT1 inhibitor reduced ATP6V1A expression in tumor lysates (**Fig. S1C**) and resulted in significantly larger mammary tumors compared to the vehicle-control treated animals (**Fig. S1D**). Moreover, exosomes isolated from their pooled serum as detected using the exosomal markers CD9 (16) and Flotillin-2 (12) (**Fig. S1E**) were enriched in ubiquitinated proteins, compared to exosomes obtained from the serum of control mice (**Fig. S1F**). Similar increases in ubiquitinated protein levels were observed in exosomes derived from the serum of mice bearing tumors of MDA-MB-231 breast cancer cells that had been depleted of SIRT1 (**Fig. S1G)**.

An examination of the Cancer RNA-Nexus database, comparing triple-negative breast cancers (TNBCs) to normal tissues adjacent to TNBC tumors, showed that more than 80% of the tumor samples exhibited a marked reduction in SIRT1 and ATP6V1A transcript levels (12), consistent with our findings from breast cancer cells and mouse models that SIRT1 down-regulation reduces ATP6V1A expression. However, this raised an important and unanswered question, namely, how does SIRT1 regulate the expression of the ATP6V1A transcript? The reduction of ATP6V1A expression due to knocking-down SIRT1 did not appear to be the result of an inhibitory effect on transcription (12). A significant clue to the underlying mechanism then emerged when we discovered that an ATP6V1A transcript which contained the coding sequence (CDS) but lacked the 3′ untranslated region (3′UTR) was not degraded under conditions where SIRT1 expression was knocked-down (**Fig. 1A**). This suggested a potential regulatory role for SIRT1 involving the 3′UTR of the ATP6V1A transcript, which is important for maintaining transcript stability. In order to pursue this lead, a biotinylated form of the 3′UTR of the ATP6V1A transcript was generated and incubated with extracts collected from SIRT1 knock-down cells (**Fig. S2A**). The biotinylated 3′UTR construct was precipitated using streptavidin-coated beads and proteins that associated with the construct were resolved by SDS-PAGE (**Fig. S2B**) and analyzed by mass spectrometry. Among the 3′UTR-associated proteins identified was IGF2BP2. *In vitro* pull-down assays using a series of truncations of the ATP6V1A 3′UTR demonstrated that IGF2BP2 binds with highest affinity to the full-length (2631 base pairs) 3′UTR construct (**Fig. 1B**, the T1 construct), while still maintaining some capability for binding to a construct consisting of only the first 824 base pairs of the 3′UTR (i.e. the T3 construct). Consistent with previous reports, we observed that IGF2BP2 has multiple translation initiation sites (17), resulting in two predominant bands being detected by Western blot analysis (Fig. 1B, lower panel).

**Figure 1.**
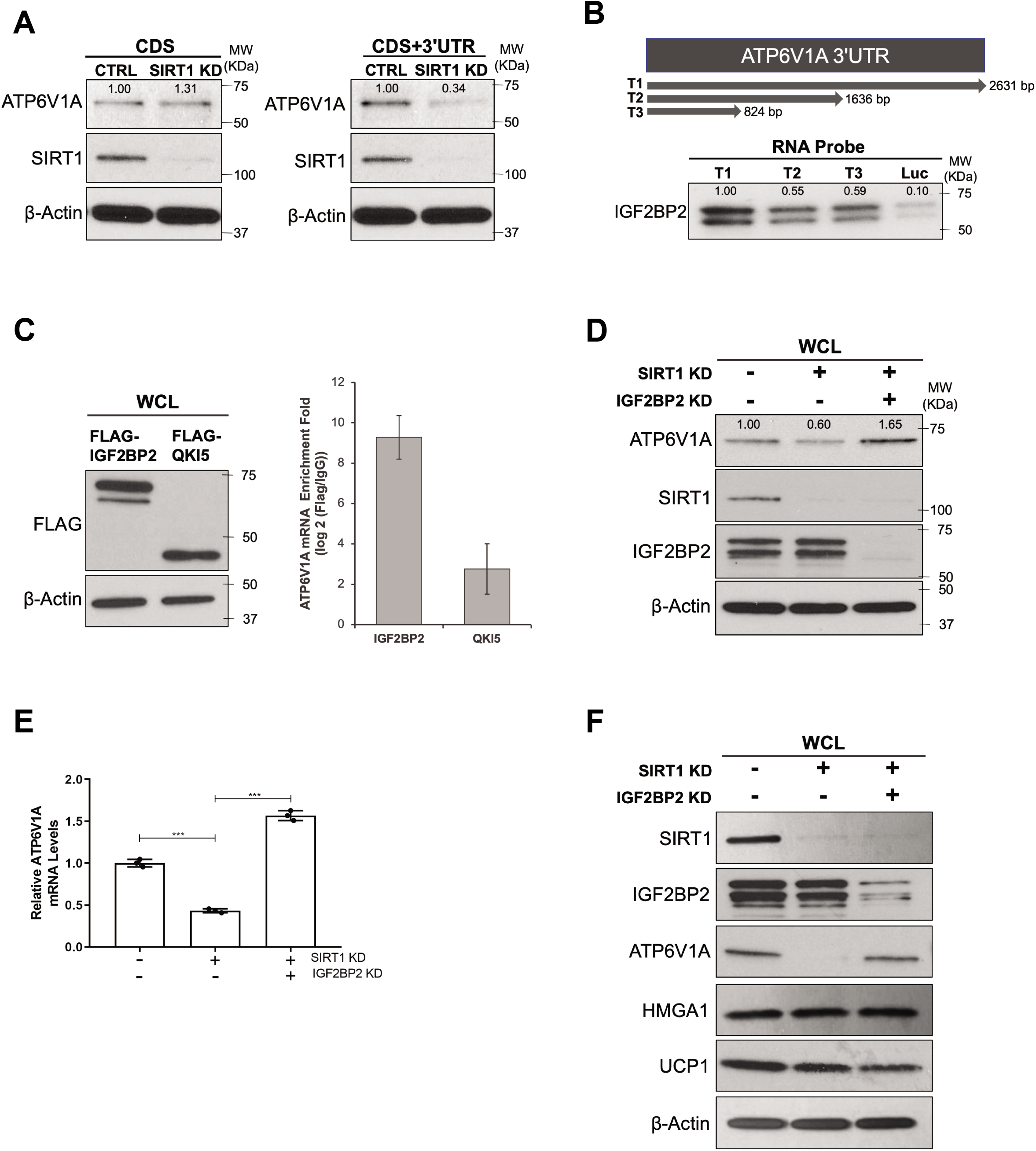
The effect of depleting cells of SIRT1 on ATP6VIA transcript levels is mediated by IGF2BP2. (A) Representative immunoblots of whole cell lysates (WCLs) from control (CTRL) and SIRT1 knock-down (SIRT1 KD) MDA-ME-231 cells ectopically expressing plasmids that encode either the coding region of the ATP6V1A (CDS), or the coding region together with its 3′UTR (CDS+3′UTR), probed for SIRT1, ATP6V1A, and β-Actin as a loading control. (B) Streptavidin pull-down assays performed on MDA-ME-231 cell WCLs incubated with the indicated biotinylated segments of the 3′UTR of the ATP6V1A transcript (diagram, top) were probed for IGF2BP2 (blot, bottom). (C) Immunoblots of WCLs from MDA-ME-231 cells ectopically expressing FLAG-tagged ATP6V1A and QKI5 probed for the FLAG-tagged proteins and β-Actin as a loading control (left). RT-qPCR was performed on the immunoprecipitated FLAG-tagged proteins to determine the levels of the ATP6V1A transcript that associated with each construct (right). (D) Representative immunoblots of WCLs from control, SIRT1 KD, and SIRT1 and IGF2BP2 KD MDA-MB-231 cells probed for ATP6V1A, SIRT1, IGF2BP2, and β-Actin as a loading control. (E) RT-qPCR was performed on the cells described in *D* to determine their expression levels of the ATP6V1A transcript (relative to actin transcript levels). The expression levels of the indicated proteins, relative to controls, in A, B, and D were quantified based on densitometry and shown above the bands. The data shown in E is presented as mean +/- SD. Statistical significance was determined using student’s t-test; *** p < 0.001. (F) Representative immunoblots of WCLs from control (-), SIRT1 KD, and SIRT1/IGF2BP2 double KD MDA-MB-231 cells probed for SIRT1, IGF2BP2, ATP6V1A, HMGA1, UCP1 and β-Actin as a loading control.

Enhanced crosslinking immunoprecipitation (eCLIP) analysis (18) highlighted potential IGF2BP2 binding sites on the ATP6V1A transcript which included a number of possible points of contact within the 3′UTR (**Fig. S2C**). When immunoprecipitations were performed from cells ectopically expressing either FLAG-tagged IGF2BP2, or FLAG-tagged Quaking 5 (QKI5), an RNA-binding protein whose homologue in *C. elegans* regulates ATP6V1A (vha-13) expression (19), we detected a significantly increased association of the ATP6V1A transcript with IGF2BP2 compared to QKI5 (**Fig. 1C**). Knocking down IGF2BP2 together with SIRT1 in cancer cells resulted in an enhancement in both the protein and transcript levels of ATP6V1A relative to control cells (**Figs. 1D**, top panel, and **1E**), demonstrating that depletion of IGF2BP2 reduced transcript degradation. These effects appear to be specific for the ATP6V1A transcript. Among the RNA transcripts that IGF2BP2 has been reported to bind (1-3) include those encoding two proteins implicated in cancer progression, High Mobility Group A1 protein (HMGA1) (20) and Uncoupling Protein 1 (UCP1) (21). However, under conditions where ATP6V1A transcript levels were reduced upon depleting SIRT1 and then restored with the knock-down of IGF2BP2, we observed only a minor reduction in UCP1 expression when knocking down SIRT1, alone or together with IGF2BP2, and no effect on HMGA1 levels (**Fig. 1F**).

Knock-down of IGF2BP2 in cells depleted of SIRT1 also caused a decrease in the number of exosomes shed (**Figs. 2A** and **2B**), and significantly reduced both the enrichment of ubiquitinated exosome cargo proteins and the amount of secreted soluble Cathepsin B (**Figs. 2C and 2D**). We then examined the effects of depleting SIRT1 and IGF2BP2 on the tumorigenicity of MDA-MB-231 cells following transplantion into the mammary fat pads of mice. Knocking down IGF2BP2 together with SIRT1 in MDA-MB-231 mouse xenografts markedly reduced tumor volumes compared to when SIRT1 was knocked-down alone (**Fig. 2E**). A significant increase in tumor mass at end point also occurred in mouse xenografts in which SIRT1 expression was knocked-down, as compared to those formed by control or SIRT1 and IGF2BP2 double knock-down cells (**Fig. 2F**).

**Figure 2.**
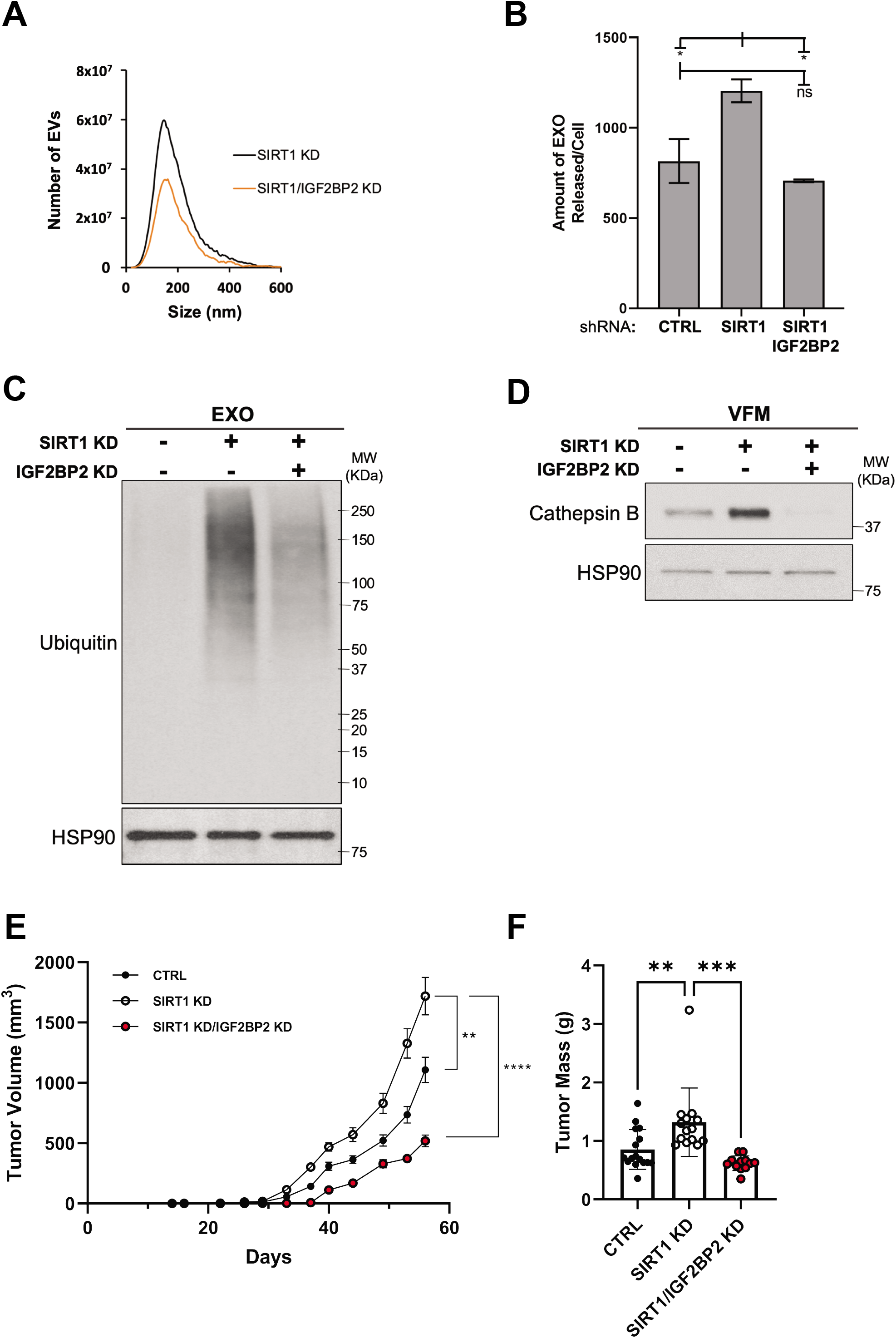
Depletion of IGF2BP2 rescues the effects caused by SIRT1 down-regulation. (A) NTA was performed on the conditioned media collected from an equivalent number of SIRT1 KD and SIRT1 and IGF2BP2 KD MDA-MB-231 cells. (B) Quantification of the assay shown in *A*. Statistical significance was determined using student’s t test; * p < 0.5, and ns, not significant. (C) Representative immunoblots of exosome lysates (EXO) prepared from control (-), SIRT1 KD, and SIRT1/ IGF2BP2 double KD MDA-MB-231 cells probed for ubiquitinated proteins and HSP90 as a loading control. (D) Representative immunoblots of vesicle free medium (VFM) prepared from the cells described in *C* probed for Cathepsin B and HSP90 as a loading control. (E and F) The volume *(E)* and mass *(F)* of the tumors that formed in mice injected with an equivalent number of control (CTL), SIRT1 KD, and SIRT1/IGF2BP2 double KD MDA-MB-231 cells were determined (n= 8, 7, and 6, respectively). The data shown in B and E represent means +/- SD. The tumor volume data represent means +/- SE at each time point, and the growth rates of the tumors were analyzed with an exponential growth model including random effects for the growth rates. Statistical significance was determined using student’s t-test; ***p < 0.001,**p < 0.01, *p < 0.05, and ns, not significant.

The ability of SIRT1 to catalyze lysine deacetylation led us to examine whether IGF2BP2 is a substrate for SIRT1 and if its acetylation was increased upon knocking down SIRT1 expression. When ectopically expressed FLAG-tagged IGF2BP2 was immunoprecipitated from SIRT1 knock-down cells and analyzed by mass spectrometry, lysine residue 530 was identified as an acetylation site (**Fig. 3A**). Western blot analysis performed on the same immunoprecipitated samples using an antibody that detects acetyl moieties showed IGF2BP2 acetylation was increased when SIRT1 expression was knocked-down (**Fig. 3B**, top panel, compare lanes 2 and 4). Changing lysine 530 to an arginine residue significantly decreased the levels of acetylated FLAG-tagged IGF2BP2 detected in SIRT1 knock-down cells (**Fig. 3B**, top panel, compare lanes 4 and 5). Moreover, ectopic expression of the acetylation-defective IGF2BP2(K530R) mutant in cells depleted of SIRT1 failed to cause a significant down-regulation of ATP6V1A expression (**Fig. 3C**, right panel), compared to cells ectopically expressing wild-type IGF2BP2 (**Fig. 3C**, left panel).

**Figure 3.**
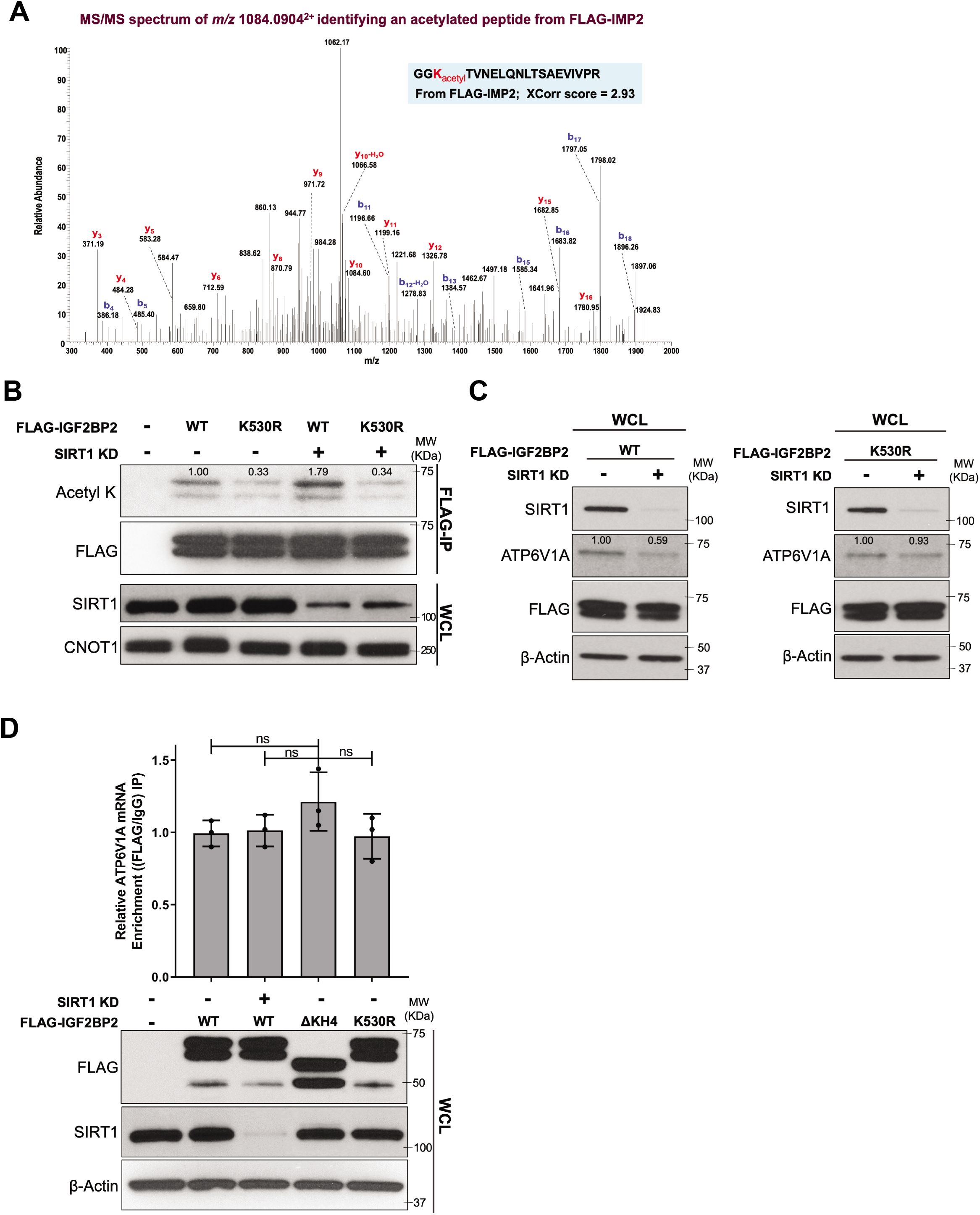
IGF2BP2 is a SIRT1 substrate. (A) Mass spectrometry profile showing the acetylated peptide identified in FLAG-tagged IGF2BP2, referred to in this figure as its alternative name IMP2, was immunoprecipitated from SIRT1 KD cells ectopically expressing this construct. The sequence, shaded in blue, represents the identified peptide and the location of the acetylated lysine (lysine 530) is highlighted (red). (B) Representative immunoblots of WCLs, as well as immunoprecipitations using a FLAG antibody (FLAG IP) performed on the WCLs, from control (-) and SIRT1 KD MDA-MB-231 cells ectopically expressing the indicated FLAG-tagged IGF2BP2 proteins probed for acetylated lysine residues (Acetyl K), SIRT1, the FLAG-tagged proteins, and CNOT1 as a loading control. (C) Immunoblots of WCLs from control (-) and SIRT1 KD MDA-MB-231 cells ectopically expressing either FLAG-tagged IGF2BP2 wild-type (WT) or the K580R mutant probed for SIRT1, ATP6V1A, the FLAG-tagged proteins, and β-Actin as a loading control. (D) Immunoblots of WCLs from control (-) and SIRT1 knockdown (SIRT 1 KD) MDA-MB-231 cells ectopically expressing the indicated forms of FLAG-tagged IGF2BP2 probed for the FLAG-tagged proteins, SIRT1, and β-Actin as a loading control (bottom). RT-qPCR was performed on the immunoprecipitated FLAG-tagged proteins to determine the levels of the ATP6V1A transcript that associated with each construct (top). The expression levels of the indicated proteins, relative to controls, in B and C were quantified based on densitometry and shown above the bands. The data shown in D is presented as mean +/- SD. Statistical significance was determined using student’s t test; ns, not significant.

Lysine 530 is located within the KH4 domain of IGF2BP2, the last of its six RNA-binding domains which includes RRM1, RRM2, and KH1-4 (**Fig. S3A**) (2). Alignment of the X-ray crystal structure of IGF2BP2 (22) with that for the highly related zip-code binding protein 1 (ZBP1) complexed to a segment of RNA (23) shows that lysine 530 is in the vicinity of a portion of the bound RNA (**Fig. S3B**). Thus, we first determined whether acetylation of lysine 530 in IGF2BP2 alters its ability to bind the ATP6V1A transcript. RNA immunoprecipitation assays were performed to compare the relative association of the ATP6V1A transcript with wild-type IGF2BP2, versus either an acetylation-defective mutant (i.e. IGF2BP2 K530R) or a truncated IGF2BP2 protein lacking the KH4 domain. However, as shown in **Fig. 3D**, neither the acetylation of lysine 530 nor the truncation of the KH4 domain affected the ability of IGF2BP2 to bind to the ATP6V1A transcript.

Since acetylation of IGF2BP2 was not an essential determinant for binding the RNA transcript encoding ATP6V1A, it seemed unlikely that IGF2BP2 alone accounted for its degradation. We therefore examined whether IGF2BP2 acetylation recruited an additional protein(s) that might be responsible for degrading the RNA transcript. Mass spectrometry was used to identify proteins that co-immunoprecipitated with IGF2BP2 in SIRT1-depleted cells. This yielded two potential IGF2BP2 binding partners reported to play roles in different aspects of RNA processing, the 5′-3′ exonuclease 2 (XRN2) and Y-box containing protein 1 (YBX1) (24,25) (**Fig. 4A**). The ability of XRN2 to bind ectopically expressed Flag-tagged IGF2BP2 as detected by their co-immunoprecipitation was significantly enhanced in SIRT1 knock-down cells (**Fig. 4B**, top panel), whereas the binding of YBX1 to IGF2BP2 was unaffected (**Fig. 4B**, second panel). Moreover, the ability of XRN2 to bind IGF2BP2 was weakened upon changing lysine 530 to arginine and eliminated when the KH4 domain was deleted (**Fig. 4C**, upper panel, top row, and **Fig. 4D**). Like the case when knocking down IGF2BP2, depleting cells of XRN2 by shRNA resulted in an increase in the transcript levels of ATP6V1A in cells lacking SIRT1 (**Fig. 4E**). The effects were specific to depleting XRN2; for example, they were not observed when knocking down CCR4-NOT transcription complex subunit 1 (CNOT1), the cytosolic de-adenylation machinery that has been shown to promote mRNA destabilization (26). Knocking down XRN2 also reduced the number of exosomes shed by SIRT1-depleted cells (**Figs. 4F and S4A**), as well as the amount of ubiquitinated proteins found in their exosome cargo (**Fig. 4G**), and significantly decreased the levels of secreted Cathepsin B (**Fig. S4B**).

**Figure 4.**
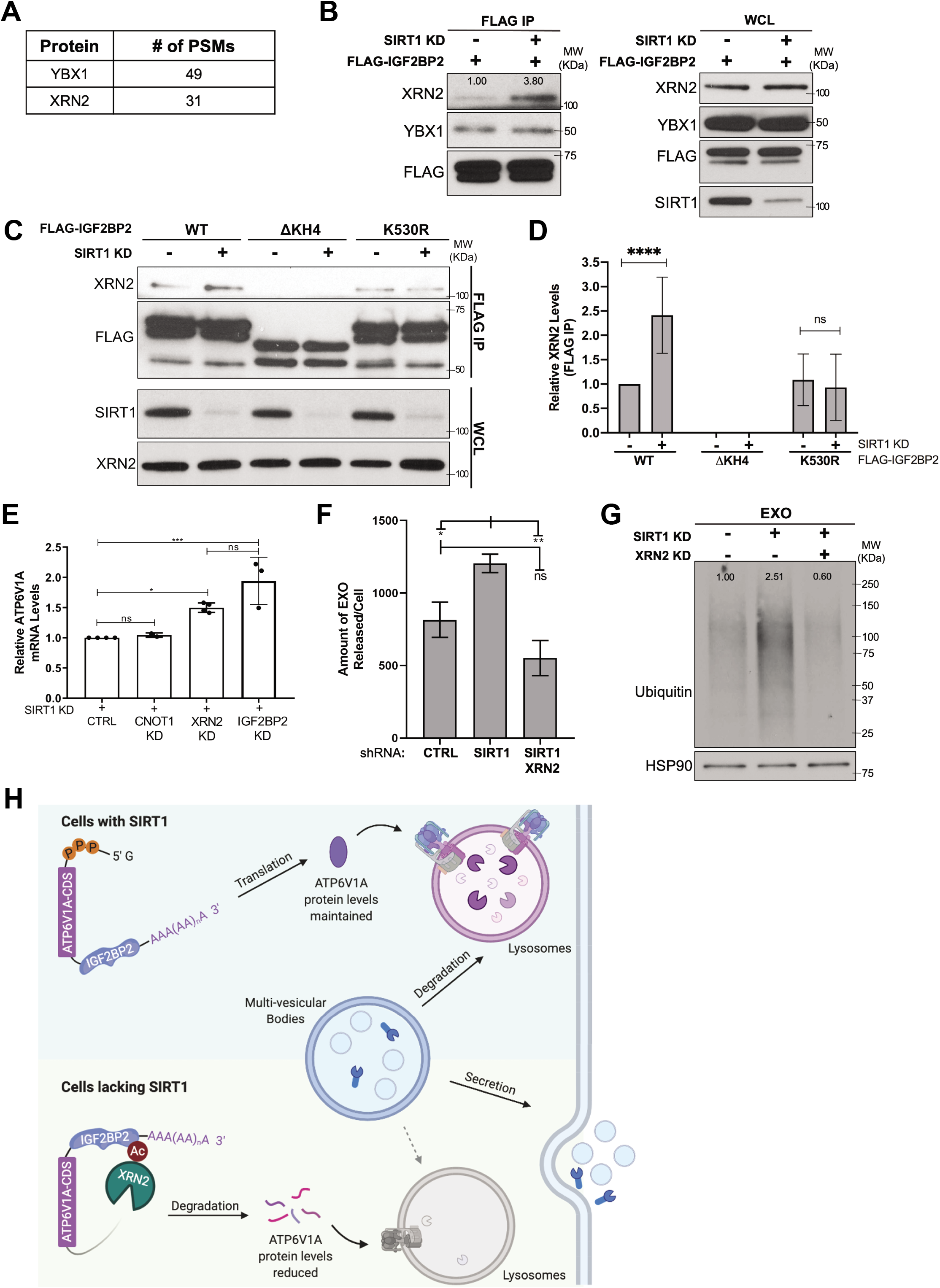
XRN2 associates with the acetylated form of IGF2BP2 and promotes ATP6V1A transcript degradation. (A) Table showing two proteins that regulate RNA stability and were found to interact with IGF2BP2 by mass spectrometry. The number of peptide spectrum matches (# of PSMs) identified for each protein are listed. (B) Representative immunoblots of WCLs, as well as immunoprecipitations using a FLAG antibody (FLAG IP) performed on the WCLs, from control (-) and SIRT1 KD cells ectopically expressing FLAG-tagged IGF2BP2 and probed for XRN2, YBX1, the FLAG-tagged construct, and SIRT1. (C) Representative immunoblots of WCLs, as well as immunoprecipitations using a FLAG antibody (FLAG IP) performed on the WCLs, from control (-) and SIRT1 KD cells ectopically expressing the indicated FLAG-tagged forms of IGF2BP2 probed for XRN2, the FLAG-tagged proteins, and SIRT1. (D) Quantification of the relative amount of XRN2 that immunoprecipitated with each of the FLAG-tagged IGF2BP2 constructs shown in *C*. (E) RT-qPCR was performed on SIRT1 KD cells (control; CTRL), or on SIRT1 KD and either CNOT1 KD, XRN2 KD, or IGF2BP2 KD cells, to determine their expression levels of the ATP6V1A transcript (relative to actin transcript levels). (F) Quantification of the NTA shown in Supplemental Figure 4A performed on the conditioned media collected from an equivalent number of SIRT1 KD, and SIRT1/XRN2 double KD cells. (G) Representative immunoblots of exosome lysates (EX0) prepared from control (-), SIRT1 KD, and SIRT1/XRN2 double KD cells probed for ubiquitinated proteins and HSP90 as a loading control. (H) Schematic diagram depicting how SIRT1 influences ATP6V1A transcript stability. In cells expressing sufficient amounts of SIRT1, the non-acetylated form of IGF2BP2 predominates and, upon binding to the ATP6V1A transcript, helps maintain its stability and thus supports lysosomal function (top). However, under conditions of limiting amounts of SIRT1, as occurs in aggressive breast cancer cells, the acetylated form of IGF2BP2 predominates, and upon binding to the ATP6V1A transcript, recruits the exonuclease XRN2. This promotes the degradation of ATP6V1A transcript, resulting in lysosomal impairment and the production of a secretome containing exosomes and hydrolases that promote aggressive phenotypes. The expression levels of the indicated proteins, relative to controls, in B and G were quantified based on densitometry and shown above the bands. The data shown in D, E, and F represent means +/- SD. Statistical significance was determined using student’s t-test; ****p < 0.0001, *** p < 0.001 **p < 0.01, *p < 0.05; ns, not significant.

In conclusion, we have discovered that the regulation of the RNA-binding protein IGF2BP2, by acting as a substrate for the NAD^+^-dependent deacetylase SIRT1, has an important role in maintaining lysosomal function. Previous studies have suggested IGF2BP2 may function in various biological and disease contexts including different forms of cancer by acting to stabilize RNA transcripts and modified long non-coding RNAs (1-8). Here we show the loss of a regulatory deacetylation of IGF2BP2, when depleting cells of SIRT1 activity, results in IGF2BP2 promoting the *degradation* of the RNA transcript that encodes ATP6V1A, a major catalytic subunit of the v-ATPase. These findings shed further light on how SIRT1 down-regulation contributes to tumor progression (12), by highlighting its unexpected role in RNA metabolism, as well as provide a mechanistic basis by which IGF2BP2 serves as a tumor promoter in breast cancer. Thus, when SIRT1 levels are down-regulated, the acetylated form of IGF2BP2 accumulates and recruits the exonuclease XRN2, thereby promoting the degradation of the ATP6V1A transcript (**Fig. 4H**). However, under conditions where SIRT1 levels are maintained, IGF2BP2 exists predominantly in a deacetylated state which is unable to recruit XRN2, ensuring the stability of the ATP6V1A transcript and proper lysosomal function. Additional factors aside from IGF2BP2 and XRN2 may be involved in providing the finely tuned regulation of ATP6V1A expression and v-ATPase function by SIRT1 since the 5′-3′ exonuclease activity of XRN2 would be expected to require decapping of the RNA transcript (27). Thus far, we have not identified an enzyme that might fulfill this function, which may be due to the transient nature of the interaction between the de-capping enzyme and the 5′ termini of mRNAs. Indeed, a similar challenge has been encountered in attempting to identify the de-capping enzyme for the case where XRN1, the cytosolic counterpart of XRN2 (28), recruits the RNA binding protein YTH domain-containing 2 (YTHDC2) to promote the degradation of YTHDC2-bound transcripts (29).

In addition to cancer progression, the newly identified roles for IGF2BP2 and SIRT1 in lysosomal maintenance may be important in other areas of biology where lysosomal impairment has been demonstrated to have significant consequences, including aging, protein homeostasis and immunity. For example, it has been reported that during aging, the levels of SIRT1and its essential co-factor NAD^+^ are reduced, and ectopic SIRT1 expression extends the lifespan of model organisms (30-34). Moreover, mice lacking both copies of IGF2BP2 exhibit longer lifespans, while caloric restriction, a lifespan-extending intervention, causes significantly reduced levels of IGF2BP2 in liver (35). Homologues of ATP6V1A are reported to be down-regulated in aging yeast and *C-elegans* (36-38), and recently it was shown that reduced levels of SIRT1 in senescent stromal cells are accompanied by decreased ATP6V1A expression and the shedding of exosomes that promote the aggressiveness of recipient cancer cells (39), and confer young cells with a senescent phenotype (40). Therefore, it will be interesting to see whether enhancing lysosomal function by promoting SIRT1 activation and IGF2BP2 deacetylation might slow down cellular senescence and aging as well as alleviate age-related neurodegenerative disorders, processes where both lysosomal impairment and increased exosome production have been implicated (41-44).

### Experimental Model and Subject Details

#### Cell Lines

Human embryonic kidney (HEK)-293T cells and MDA-MB-231 cells were obtained from the ATCC (https://www.atcc.org). The MCF10A, MCF10AT1, and MCF10CA1A cells were kindly provided by Claudia Fischbach, Cornell University. HEK-293T cells were cultured in Dulbecco’s Modified Eagle Medium (DMEM) supplemented with 10% calf serum (CS), while MDA-MB-231, MCF10A, MCF10AT1, and MCF10CA1A cells were cultured in Roswell Park Memorial Institute 1640 (RPMI-1640) medium supplemented with 10% fetal bovine serum (FBS). Cells stably expressing the indicated constructs were selected for, and maintained by, supplementing the growth medium with 2.0 μg/mL puromycin. All cell lines were maintained at 37°C with 5% CO2.

#### Animal Model

All mice used in this study were handled following federal and institutional guidelines under a protocol approved by the Institutional Animal Care and Use Committee (IACUC) at Cornell University. Starting at 4 weeks of age, MMTV-PyMT transgenic female mice were palpated every other day for mammary tumors. Upon the detection of tumors, mice were treated with 14 mg/kg EX-527 (dissolved in 70% PEG-300, 5% DMSO and 25% ddH_2_O), or the vehicle alone, by intraperitoneal (IP) injections every other day for 5 weeks. Control, SIRT1 KD, and SIRT1/IGF2BP2 double KD MDA-MB-231 cells (3.0 × 10^6^) were injected into the mammary fat pad of 8-week old female NSG mice. The health of the animals was monitored daily, and their weight was determined every other day. Tumor size was measured using calipers every other day, and tumor volumes were calculated using the formula 0.5 x (length x width^2^). Either at the completion of the experiment, or when the mice reached a humane endpoint, they were euthanized using CO2 asphyxiation, and the tumors were collected, weighed, lysed using lysis buffer (25 mM Tris, 100 mM NaCl, 1.0 % Triton X-100, 1.0 mM EDTA, 1.0 mM DTT, 1.0 mM NaVO4, 1.0 mM β-glycerol phosphate, and 1.0 μg/ml each aprotinin and leupeptin), and stored at -80°C. Serum was also collected from the mice (see below).

### Method Details

#### Plasmid Generation and Site-directed Mutagenesis

The lentiviral pLJM1 expression plasmid containing a C-terminal FLAG tag was obtained from Addgene (#91980), and the genes encoding IGF2BP2 and QKI5 were cloned into the plasmid using the InFusion Cloning Kit (Clonetech) and the primers CGTCAGATCCGCTAGCATGATGAACAAGCTTTACATCGGG and TCGAGGTCGAGAATTCTCACTTGTCGTCATCGTCTTTGTAGTCACTACCTCCACCTCC CTTGCTGCGCTGTGAG for IGF2BP2, and CGTCAGATCCGCTAGCATGGTCGGGGAAATGGAAACG and TCGAGGTCGAGAATTCTTACTTGTCGTCATCGTCTTTGTAGTCACTACCTCCACCTCC GTTGCCGGTGGCGGC for QKI5. Mutation of lysine (K) 530 to an arginine (R), as well as deletion of the KH4 domain, in IGF2BP2 was carried-out using the InFusion Cloning Kit (Clonetech) and the primers: AGGTGGCAGGACCGTGAACGAACTGCAGAAC and ACGGTCCTGCCACCTTTGCCAATCACCC for generating the IGF2BP2-K530R mutant, and AAGAGGAAGGAGGTGGAGGTAGTGAC and CACCTCCTTCCTCTTTCAGTTTCCCAAAG for generating IGF2BP2-ΔKH4 mutant.

#### Virus Production and Cell Infection

Lentiviruses were generated by transfecting HEK-293T cells with the pLJMl expression plasmids, a sham-targeting (control) shRNA plasmid (Sigma), or with shRNAs that target SIRTl, IGF2BP2, XRN2, and CNOTl, and the packaging plasmids (#12259 and #12263, Addgene) using Fugene 6 (Promega). The viruses shed into the medium by the cells were harvested 24 and 48 h after the transfection. To infect cells, they were treated with the various viruses generated and Polybrene (8 μg/mL).

#### Exosome Isolation from Conditioned Medium, Vesicle Free Medium (VFM) Preparation, and Whole Cell Lysate Generation

The conditioned medium collected from 2.0 × 10^6^ serum starved cells was subjected to two consecutive centrifugations at 700 x g to clarify the medium of cells and debris. The partially clarified medium was filtered using a 0.22 μm pore size Steriflip PVDF filter (Millipore), and the filtrate was subjected to ultracentrifugation at 100,000 x g for 8 h. The exosome pellets were then either lysed in 250 μL of lysis buffer, or resuspended in 500 μL of sterile phosphate buffered saline (PBS). The medium depleted of MVs and exosomes was then concentrated using Centricons with a 10 KDa pore size (Amicon), and was considered as the vesicle-free medium (VFM) samples. Whole cell lysates (WCL) were prepared by rinsing dishes of cells with PBS, adding 800 μL of lysis buffer, and scraping the cells off of the plate. The exosome lysates, WCLs, and VFM preparations were centrifuged at 16,000 x g for 10 min, and the supernatants were collected and stored at -80°C.

#### Collection of EVs from Serum

Approximately 500 μL of blood was collected from each tumor-bearing MMTV-PyMT transgenic female mouse that had been treated with EX-527, or the vehicle control. Similar amounts of blood were also collected from NSG mice that had been injected with control or SIRT1 KD MDA-MB-231 cells. The samples were centrifuged at 3,000 x g for 10 min, and the resulting serum fraction collected from 3 similarly treated mice were pooled and then diluted in 2.0 mL PBS. The samples were subjected to two consecutive centrifugations at 700 x g to clarify them of cells and debris, before being centrifuged at 100,000 x g at 4°C for 8 h. The pellets were washed with 3.0 mL PBS and centrifuged again at 100,000 x g at 4°C for another 4 h. The exosome pellet was lysed in 250 μL of lysis buffer, centrifuged at 16,000 x g for 10 min, and the supernatant was collected and stored at -80°C

#### Immunoprecipitations

Equal concentrations (1.0 mg) of WCLs from MDA-MB-231 cells ectopically expressing the indicated FLAG-tagged forms of IGF2BP2 were incubated with FLAG antibody labeled magnetic beads (Sigma) according to the manufacturer’s instructions. The beads were precipitated using a magnetic rack, and then were resuspended in 100 μL of 150 ng/μL of 3X-FLAG peptide (Sigma) diluted in TBS buffer (50 mM Tris-Cl (pH 7.5), 150 mM NaCl). The beads were precipitated again using the magnetic rack, and the supernatants were collected and stored at -80°C. For the experiments where changes in the levels of FLAG-tagged forms of IGF2BP2 were determined, the WCLs were supplemented with 10 mg/mL each of nicotinamide and sodium butyrate to prevent deacetylation.

#### Immunoblot Analysis

The protein concentrations of WCLs, exosome lysates, and VFM preparations were determined using the Bradford protein assay and normalized by protein concentration. These samples, as well as the reactions from the protease protection assays and immunoprecipitations performed as described above, were resolved by SDS-PAGE and the proteins were transferred to PVDF membranes. The membranes were blocked with 5% bovine serum albumin (Sigma) in TBST (19 mM Tris Base, 2.7 mM KCl, 137 mM NaCl, and 0.5 % Tween-20) for 1 h, and then they were incubated with the indicated primary antibodies. The next day, the primary antibodies were detected using HRP-conjugated secondary antibodies (Cell Signaling Technology) and exposure to ECL reagent (Pierce).

#### Mass Spectrometry

FLAG-tagged IGF2BP2 that had been immunoprecipitated from the indicated WCLs using a FLAG antibody, was used to identity the lysine residues that were acetylated in this protein, as well as determine proteins that associated with IGF2BP2, using LC-MS/MS (Proteomics and Metabolomics Facility, Cornell University).

#### Nanoparticle Tracking Analysis

The sizes and numbers of EVs in conditioned medium collected from an equal number of serum starved cells were determined using the NanoSight NS300 (Malvern, Cornell NanoScale Science and Technology Facility) as described in Kreger et al., 2016. Briefly, the samples were injected into the beam path to capture movies of EVs as points of diffracted light moving rapidly under Brownian motion. Five 60-s digital videos of each sample were taken and analyzed to determine the sizes and numbers of EVs based on their movement, and then results were averaged together.

#### In vitro RNA Transcription

The T7 promoter was added to the full length cDNA encoding the 3′UTR of the ATP6V1A transcript (referred to as T1), and the indicated truncations of this 3′UTR (referred to as T2 and T3), by performing PCR and using the ATP6V1A cDNA containing its 3′UTR as a template, the T7 promoter primer CTAATACGACTCACTATAGGGAGAAAGCCTTGAAGATTACAACTG, and one of the following primers: TGTTAATTTAAATCCACTTTTTATTCTTTCACAG to generate the Tl construct, CAGAGCTGTTCTGCAATATGCAGACAC to generate the T2 construct, or TGACCAATATGGTGAAACCCCGTTTCTAC to generate the T3 construct. *In vitro* transcription of the Tl, T2, and T3 constructs were performed using the MEGAscript T7 Transcription Kit (New England Bioenzymes) according to the manufacturers’ instructions, and the resulting transcripts were biotinylated using the Pierce™ RNA 3′ End Biotinylation Kit (Thermo Fisher).

#### Biotin Pull-down Assays

The biotinylated RNA transcripts representing different portions of the ATP6V1A 3′UTR, i.e. the T1-T3 constructs, (100 pmol) were incubated for 1 h at room temperature with magnetic streptavidin-conjugated beads (New England Bioenzymes, 100 uL per sample) that had been prepared according to the manufacturer’s instructions. The beads were then collected using a magnetic rack, washed twice with 400 μL of binding and wash buffer (5 mM Tris-HCl (pH7.4), and 1 M NaCl), and were resuspended in 50 μL of buffer C (25 mM Tris-HCl (pH 7.4), 150 mM NaCl, 2 mM MgCl_2_, 1 mM DTT, 1 mM PMSF, 1X protease inhibitor cocktail, and 0.4 U/μl RNase inhibitor). WCLs (750 μg of each) were then incubated with the RNA-conjugated beads for 3 h at 4 °C with rotation. The beads were then precipitated using a magnetic rack, washed six times with 1.0 mL of buffer C containing 40 U RNase inhibitor and 0.25% IGEPAL, resuspended in 75 μL of buffer C and SDS-PAGE sample buffer, and boiled for 5 min. The samples were centrifuged at 12,000 x g for 1 min and the supernatant was resolved by SDS-PAGE and either silver-stained or subjected to Western blot analysis.

#### RNA Immunoprecipitation (RIP)

To immunoprecipitate RNA from WCLs, the Imprint® RNA Immunoprecipitation Kit (Sigma) was used according to the manufacturer’s instructions. Briefly, WCLs of MDA-MB-231 cells ectopically expressing the indicated FLAG-tagged forms of IGF2BP2 (1.0 mg of each) were incubated with either 1.0 μg of FLAG antibody (Cell Signaling Technologies), or 1.0 μg of mouse isotype control antibody (Cell Signaling Technologies). Following the immunoprecipitation of the FLAG-tagged forms of IGF2BP2 as described above, the RNA transcripts that associated with the ectopically expressed proteins were isolated using the Direct-zol RNA Microprep Kit (Zymo Research).

#### RNA Isolation and Quantitative (q) RT-PCR Analysis

Total RNA was isolated from cells using the Direct-zol RNA Microprep Kit (Zymo Research), and the mRNA transcripts were converted to cDNA using Superscript III Reverse Transcriptase (Invitrogen) and oligo dT2o. The cDNA was then used to determine the ATP6V1A and β-actin (as a control) transcript levels using SYBR Green Supermix (Bio-Rad), the Applied Biosystems® 7500 Real-Time PCR System with the T method (ABI), and the primers ACATCCCCAGAGGAGTAAACG and ACTACCAACCCGTAGGTTTTTG for amplifying ATP6V1A, and CATGTACGTTGCTATCCAGGC and CTCCTTAATGTCACGCACGAT for amplifying β-actin.

#### Quantification and Statistical Analysis

Quantitative data are presented as means ± SD. All experiments were independently performed at least three times. Statistical significance was calculated by ANOVA (Tukey’s test) for experiments involving comparing more than two conditions, and student’s t test for experiments involving comparing two conditions. Error bars represents the mean ± SD. *p ≤ 0.05, **p ≤ 0.01, ***p<0.001, ****p<0.0001, n.s. = non-significant.

## Acknowledgments

This research was supported by grants from the NIH <u>(R35GM122575</u> and <u>R01CA201402</u>) to R.A.C., (U54CA210184) to C.F., (R01 CA223534) to R.A.C., H.L., and R.S.W., <u>(DK107868)</u> to H.L., (F30 CA25045) to J.J.M, grants from the NIH (T32GM007273), a Cornell Deans Excellence Fellowship, and a HHMI Gilliam Fellowship to I.R.F., and grants from the NIH <u>(F99 CA234921</u>) and The Breast Cancer Coalition of Rochester to A.L. NTA was performed at the Cornell NanoScale Facility and was supported by NSF grant <u>NNCI-1542081</u>.

## Author Contributions

A.L., F.W., J.M., I.F. and L.L. performed the experiments. A.L., F.W., C.F., H.L., R.S.W, R.A.C., and M.A.A. designed the project and wrote the manuscript.

## Declaration of Interests

The authors declare no competing interests.

## Supplemental Figure Legends

**Supplemental Figure 1. SIRT1 downregulation produces a secretome that promotes breast cancer growth**.

(A) Representative immunoblots of WCLs from MCF10A, MCF10AT1, and MCF10CA1A cells probed for SIRT1, ATP6V1A, and β-Actin as a loading control. (B) Nanoparticle tracking analysis was performed on the conditioned medium collected from an equivalent number of MCF10A and MCF10CA1A cells shown in *A*. (C) Representative immunoblots of tumor lysates collected from tumor-bearing PYMT mice treated every other day for 5 weeks with vehicle or EX-527 (14 mg/Kg) probed for ATP6V1A and β-Actin as a loading control. (D) Total tumor mass (top) and images of tumors from representative individual mice (bottom) collected following treatment as described in *C* (n= 10 and 6 for mice treated with vehicle and EX-527, respectively). (E) Representative immunoblots of tumor and extracellular vesicle lysates (EV) prepared from the serum of tumor-bearing PYMT mice probed for CD9, Flottilin-2, and IKBα. (F) Immunoblots of extracellular vesicle lysates (EV) prepared from the serum of tumor-bearing PYMT mice treated without (Vehicle) or EX-527 probed for ubiquitinated proteins and Flotillin-2 as a loading control. (G) Representative immunoblots of extracellular vesicle lysates prepared from the serum of tumor-bearing control (CTRL) and SIRT1 KD MDA-MB-231 mouse xenografts. The expression levels of the indicated proteins, relative to controls, in A and C were quantified based on densitometry and shown above the bands. The data shown in D is presented as mean +/- SD. Statistical significance was determined using student’s t-test; **p < 0.01.

**Supplemental Figure 2. Identification of proteins that associate with the 3′UTR of the ATP6V1A transcript**.

(A) Diagram showing the approach used to identify proteins that associate with the 3′UTR of the ATP6V1A transcript. (B) An SDS-PAGE gel of streptavidin pulldowns performed on SIRT1 KD MDA-MB-231 cells incubated with either beads alone (Beads) or a biotinylated form of the 3′UTR of the ATP6V1A transcript (3′UTR). The gel was stained with Colloidal Blue to label proteins. (C) eCLIP analysis was performed on either a mock RNA transcript (Mock Input eCLIP, top), or the full length ATP6VIA RNA transcript (IGF2BP2 eCLIP, bottom). The location of introns, exons, and the 3′UTR in the ATP6V1A transcript are indicated, and the 3′UTR is further boxed in red.

**Supplemental Figure 3. The acetylated lysine in IGF2BP2 is located in an RNA binding domain**.

(A) Linear depiction of the six different RNA binding domains in IGF2BP2. The acetylated lysine in this protein identified by mass spectrometry is residue 530, which is located in the GXXGL loop (highlighted in red) of its KH4 domain. (E) Alignment of a portion of the X-ray structure of the IGF2BP2 KH4 domain (PDE: 2N8M) with the X-ray structure for the corresponding region of ZBP1 bound to RNA (24). The positioning of the GXXGL loop (cyan), lysine residue 530 (magenta), and the RNA molecule (orange) are all shown.

**Supplemental Figure 4. Mouse models showing the effects of depleting cancer cells of SIRT1, IGF2BB2, and XRN2 on tumor mass**.

(A) NTA was performed on the conditioned media collected from an equivalent number of SIRT1 KD and SIRT1/XRN2 double KD MDA-ME-231 cells. This assay was quantification in Figure 4F. (E) Representative immunoblots of the vesicle-free media (VFM) collected from control (-), SIRT1 KD, and SIRT1/XRN2 double KD cells probed for Cathepsin B and HSP90 as a loading control. Quantification based on densitometry is shown above the bands in the top panel.

## Notes

### Competing Interest Statement

The authors have declared no competing interest.

