## Supplementary figures and images for "IGF2BP2 Promotes Cancer Progression by Degrading the RNA Transcript Encoding a v-ATPase Subunit"

### Supplemental Figures

#### Supplemental Figure 1

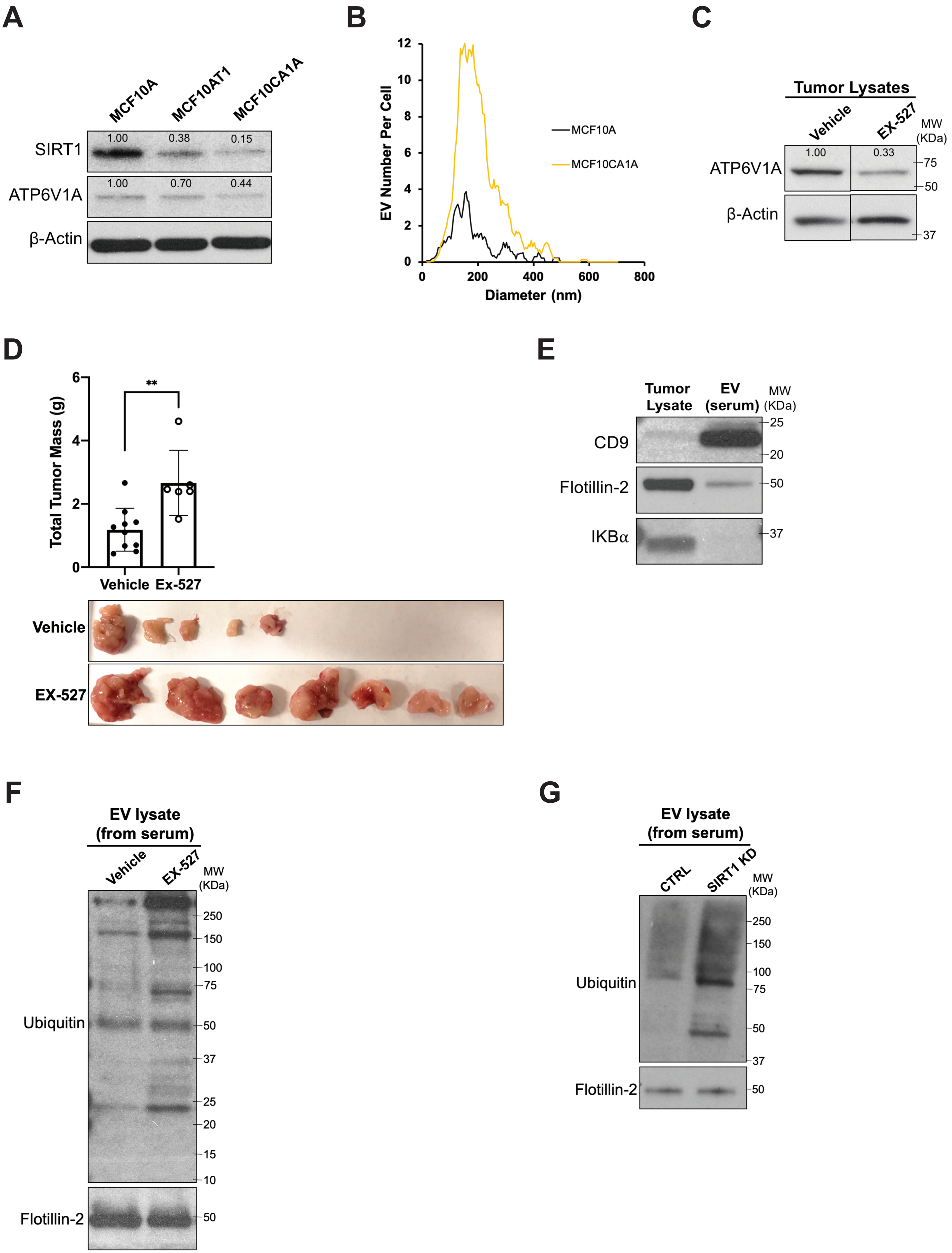

#### Supplemental Figure 2

**A**

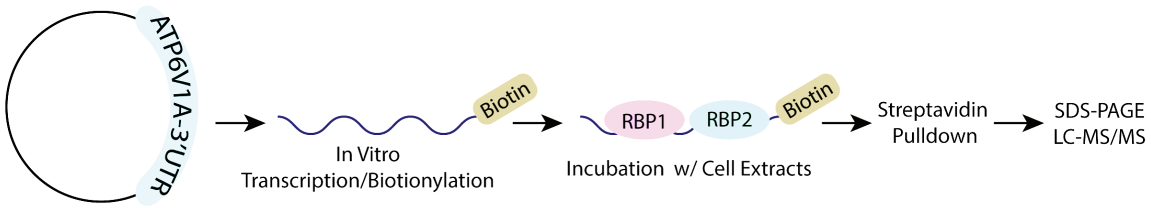


**B**

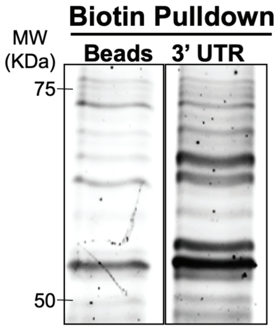


**C**

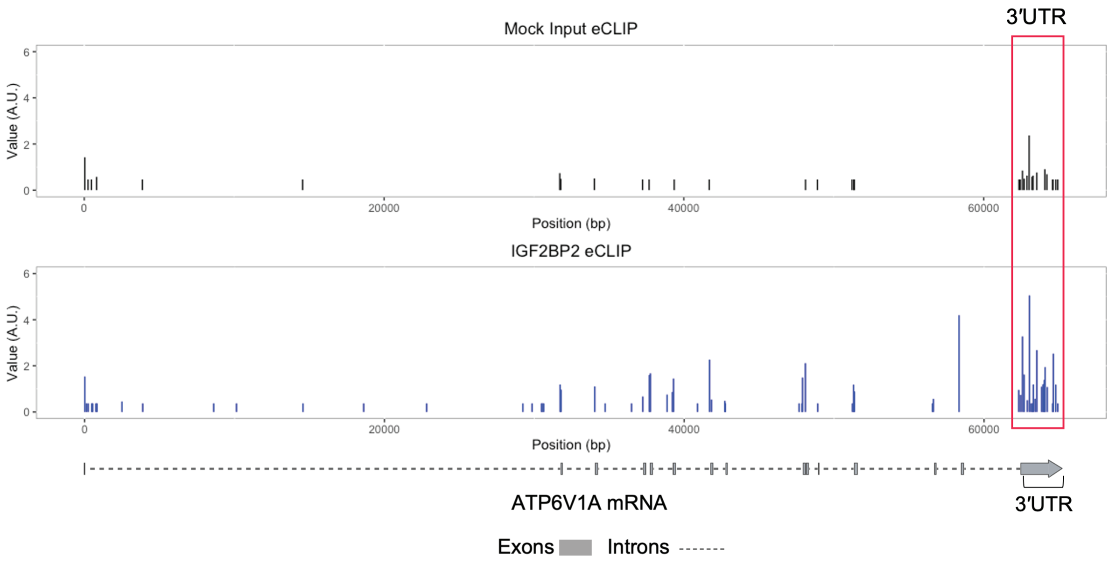

#### Supplemental Figure 3

**A**

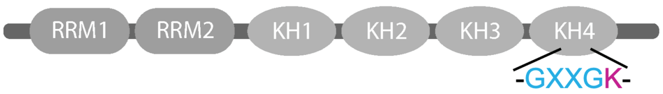


**B**

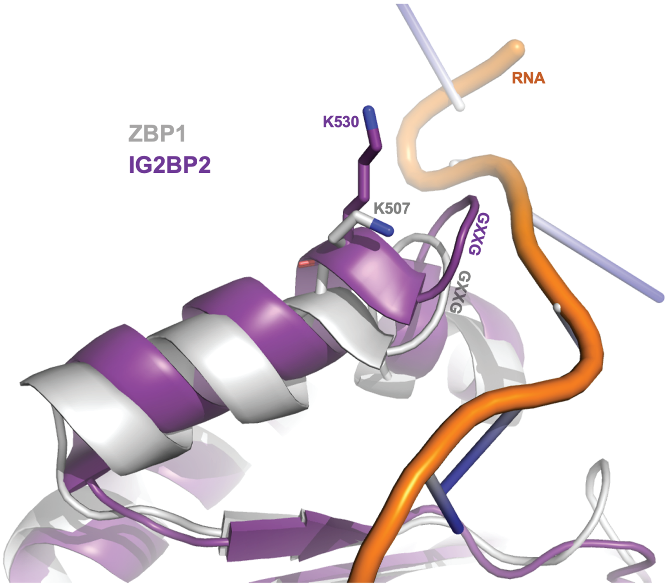

#### Supplemental Figure 4

**A**

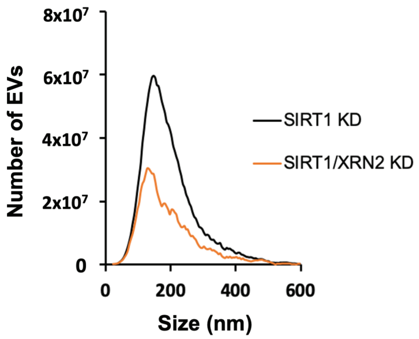


**B**

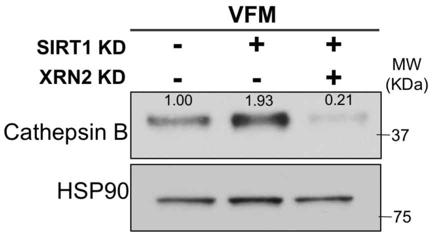
